# The fission yeast cytokinetic ring component Fic1 promotes septum formation

**DOI:** 10.1101/2023.03.13.532462

**Authors:** Anthony M. Rossi, K. Adam Bohnert, Kathleen L. Gould

## Abstract

In *Schizosaccharomyces pombe* septum formation is coordinated with cytokinetic ring constriction but the mechanisms linking these events are unclear. In this study, we explored the role of the cytokinetic ring component Fic1, first identified by its interaction with the F-BAR protein Cdc15, in septum formation. We found that the *fic1* phospho-ablating mutant, *fic1-2A*, is a gain-of-function allele that suppresses *myo2-E1*, the temperature-sensitive allele of the essential type-II myosin, *myo2*. This suppression is achieved by the promotion of septum formation and required Fic1’s interaction with the F-BAR proteins Cdc15 and Imp2. Additionally, we found that Fic1 interacts with Cyk3 and that this interaction was likewise required for Fic1’s role in septum formation. Fic1, Cdc15, Imp2, and Cyk3 are the orthologs of the *Saccharomyces cerevisiae* ingression progression complex, which stimulates the chitin synthase Chs2 to promote primary septum formation. However, our findings indicate that Fic1 promotes septum formation and cell abscission independently of the *S. pombe* Chs2 ortholog. Thus, while similar complexes exist in the two yeasts that each promote septation, they appear to have different downstream effectors.

**Summary Statement:** The *S. pombe* cytokinetic ring protein Fic1 promotes septum formation in a manner dependent on interactions with the cytokinetic ring components Cdc15, Imp2, and Cyk3.

## Introduction

Cytokinesis is the final process in the cell cycle which creates two independent daughter cells. Many eukaryotic organisms use an actin-myosin structure known as the cytokinetic ring (CR) to mark the plane of cell division and to drive membrane ingression (reviewed in (Cheffings et al., 2016; Mangione and Gould, 2019)). In organisms with cell walls, such as *Schizosaccharomyces pombe* and *Saccharomyces cerevisiae*, the CR alone is insufficient for cytokinesis (Jochova et al., 1991; Munoz et al., 2013; Proctor et al., 2012; Ramos et al., 2019; Schmidt et al., 2002). These organisms require the formation of a septum coupled to CR constriction to drive cell abscission (Cortes et al., 2007; Cortes et al., 2015; Jochova et al., 1991; Proctor et al., 2012; Schmidt et al., 2002).

Yeast septa are trilaminar structures composed of a primary septum flanked by secondary septa (Humbel et al., 2001; Wloka and Bi, 2012). In *S. pombe* and *S. cerevisiae*, CR constriction promotes septation perpendicular to the cell cortex (Cortes et al., 2002; Cortes et al., 2007; Johnson et al., 1973; Roncero et al., 2016; Schmidt et al., 2002). In *S. cerevisiae* the chitin synthase Chs2 polymerizes N-acetylglucosamine to form the primary septum (Sburlati and Cabib, 1986; Shaw et al., 1991; Silverman et al., 1988). In contrast, in *S. pombe* it is the glucan synthases Bgs1 and Ags1 that polymerize linear-β(1,3)glucans and α(1,3)glucans, respectively, to form the primary septum (Cortes et al., 2002; Cortes et al., 2007; Cortes et al., 2015; Cortes et al., 2012).

*S. cerevisiae* Chs2 and septum formation are stimulated by a protein complex within the CR named the ingression progression complex (IPC), comprised of the ingression protein Inn1, the F-BAR protein Hof1, and Cyk3 (Devrekanli et al., 2012; Nishihama et al., 2009; Sanchez-Diaz et al., 2008). Analogous proteins exist in *S. pombe*. Specifically, *S. pombe* Fic1, Cdc15/Imp2, and Cyk3 are the orthologs of Inn1, Hof1, and Cyk3, respectively (Demeter and Sazer, 1998; Fankhauser et al., 1995; Pollard et al., 2012; Roberts-Galbraith et al., 2009). Fic1 was identified in a yeast-two hybrid screen using the SH3 domains of Cdc15 as bait and directly interacts with Cdc15 and Imp2 (Ren et al., 2015; Roberts-Galbraith et al., 2009). *S. pombe* Cyk3 was identified based on sequence similarity to *S. cerevisiae* Cyk3 and has been found to co-immunoprecipitate with Fic1 (Bohnert and Gould, 2012; Roberts-Galbraith et al., 2009). However, it is unknown if these *S. pombe* proteins cooperate to promote primary septum formation similarly to the IPC.

We previously found that Fic1 is phosphorylated on two sites by multiple kinases (Bohnert et al., 2020). Preventing phosphorylation at these sites produces defects in the establishment of normal cell polarity. Here, we pursued the observation that the *fic1* phospho-ablating mutant, *fic1-2A*, also suppressed the *myo2-E1* temperature-sensitive allele of the essential type-II myosin Myo2 (Balasubramanian et al., 1998; Kitayama et al., 1997). The inability of *myo2-E1* cells to form a functional CR to guide septum formation prevents cytokinesis and leads to cell death (Balasubramanian et al., 1998). Time-lapse microscopy showed that *fic1-2A* suppressed *myo2-E1* by promoting septum formation and daughter cell abscission and that cells lacking *fic1* exhibited significant delays in septation. We determined that the ability of *fic1-2A* to suppress *myo2-E1* required its interactions with Cyk3, Cdc15, and/or Imp2 but not Chs2. This work revealed that *S. pombe*’s IPC analogs interact to promote septum formation through a mechanism that is functionally divergent from the IPC in *S. cerevisiae*.

## Results and Discussion

### Fic1 phospho-ablating mutant suppresses *myo2-E1*

To determine if Fic1’s phosphorylation state impacts cytokinesis, we took a genetic approach and probed interactions between *fic1* phosphomutants and deletions or temperature-sensitive alleles of genes involved in actin dynamics (*cdc12*), septum formation (*sid2, bgs1*, and *bgs4*), and CR constriction (*cdc4* and *myo2*) (Fig. S1A). From this screen we observed one significant interaction: *fic1*’s phospho-ablating mutant, *fic1-2A*, suppressed *myo2-E1* (Fig. 1A,B and S1A). *myo2-E1* is a temperature-sensitive allele of the essential type-II myosin, *myo2*, that inhibits Myo2’s activity and produces non-constricting CRs at the restrictive temperature (Balasubramanian et al., 1998; Palani et al., 2017; Palani et al., 2018). Without CR constriction, cell wall accumulates at the division site but does not form a septum (Balasubramanian et al., 1998; Palani et al., 2017; Palani et al., 2018; Ramos et al., 2019). Interestingly, *fic1Δ* did not suppress *myo2-E1* and no genetic interaction was observed between *fic1-2D* and *myo2-E1* (Fig. 1B,C), which suggests *fic1-2A* is a gain-of-function allele but whether this gain in function is due to alterations to Fic1’s phosphorylation state is unclear. The individual phospho-ablating *fic1* mutants only partially suppressed *myo2-E1* and none of the *fic1* phosphomutants were temperature sensitive (Fig. S1B,C). We then pursued the underlying mechanisms behind *myo2-E1*’s suppression to gain insight into the cytokinetic roles of Fic1, a CR protein of enigmatic function.

**Figure 1.**
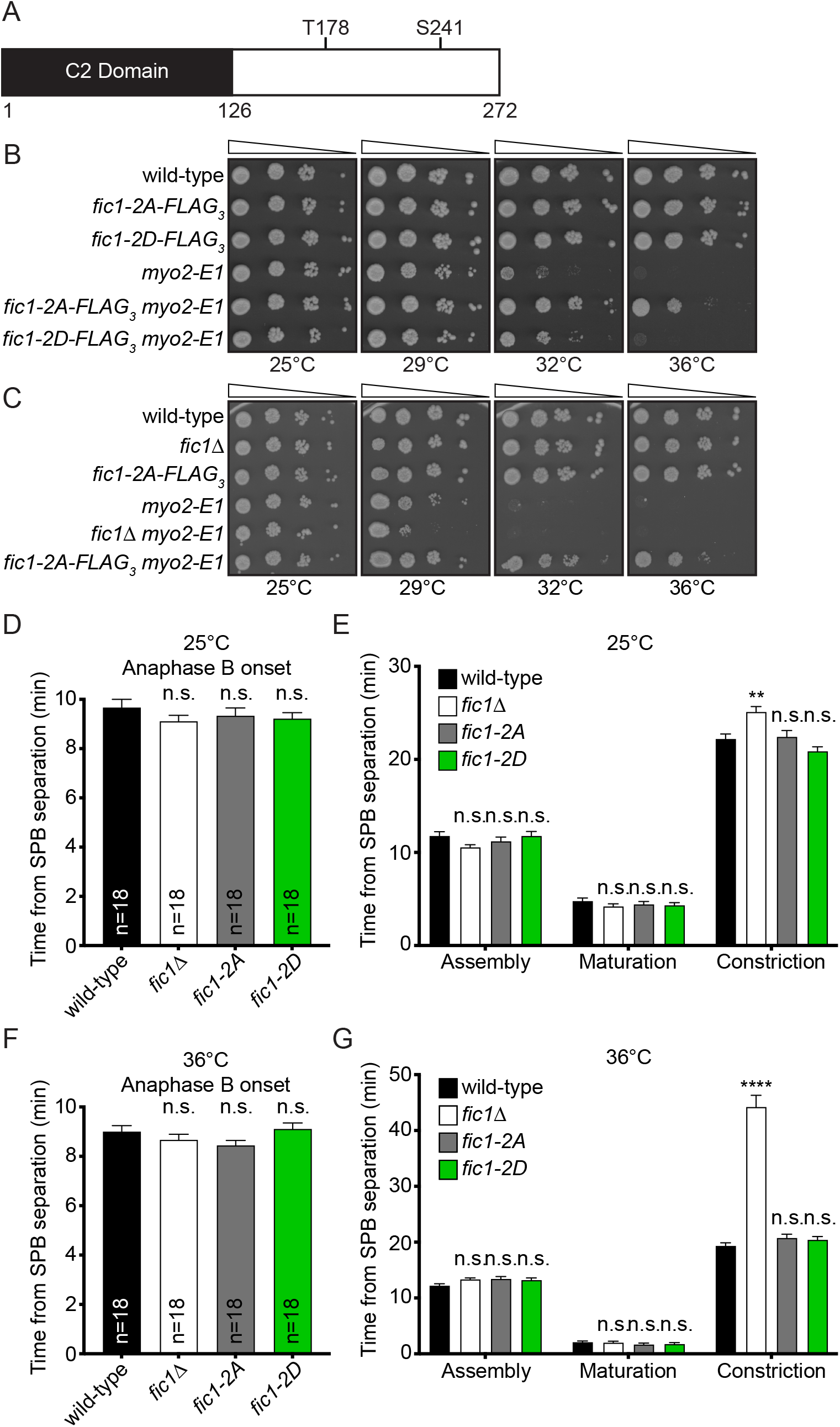
*fic1-2A* suppresses *myo2-E1*. A) Schematic of Fic1 with domain boundaries and phosphorylation sites indicated. Drawn to scale. B and C) Ten-fold serial dilutions of the indicated strains were spotted on YE agar media and incubated at the indicated temperatures for 3-5 days. D-G) Quantification of timing of anaphase B (D and F) and CR assembly, maturation, and constriction (E and G) for each strain at the indicated temperatures. n=number of cells analyzed. Data presented as mean ± S.E.M. ****p≤0.0001, **p≤0.01, n.s., not significant, one-way ANOVA.

### *fic1-2A* cells exhibit similar CR dynamics compared to wild-type cells

We postulated that *myo2-E1* suppression by *fic1-2A* could be achieved by altering CR dynamics. *fic1-2A* could provide additional time for proper glucan synthase localization by prolonging CR maturation and/or constriction. Glucan synthases are trafficked to the site of cell division and localize diffusely on the cortex (Cortes et al., 2002; Hoya et al., 2017; Katayama et al., 1999; Mulvihill et al., 2006; Ramos et al., 2019). As the CR constricts the glucan synthases coalesce into a ring concentric with the CR, Bgs1 is activated, and primary septum formation begins (Ramos et al., 2019). By providing additional time for glucan synthase ring formation by prolonging CR maturation and/or constriction *fic1-2A* could effectively promote septum formation. Alternatively, *fic1-2A* could increase the rate of CR constriction which could allow *fic1-2A* to suppress *myo2-E1* by restoring the contractile function of the CR and septum formation.

To test these possibilities, we performed live-cell time-lapse imaging at 25°C and 36°C of wild-type, *fic1Δ, fic1-2A*, and *fic1-2D* cells containing a CR marker, *rlc1-mNG*, to monitor CR dynamics and a spindle pole body (SPB) marker, *sid4-GFP*, to monitor mitotic progression (Chang and Gould, 2000; Naqvi et al., 2000). The timing of anaphase B onset was similar between all strains at both temperatures (Fig. 1D,F), as was the timing of CR assembly and CR maturation (Fig. 1E,G). The timing of CR constriction was similar between wild-type, *fic1-2A*, and *fic1-2D* cells at both temperatures indicating that Fic1 phosphostate does not appreciably affect CR dynamics (Fig. 1E,G). However, *fic1Δ* took longer, an average of 25.1±0.6 and 44.2±2.1 minutes at 25°C and 36°C, respectively whereas wild-type cells took 22.2±0.5 and 19.3±0.6 minutes at 25°C and 36°C, respectively (Fig. 1E,G). Delayed CR constriction in *fic1Δ* but not *fic1-2A* suggests that prolonging CR constriction is not how *fic1-2A* suppresses *myo2-E1*. Rather, because the rate of CR constriction is linked to the rate of septum deposition (Proctor et al., 2012; Ramos et al., 2019), the delay in CR constriction of *fic1Δ* suggests that Fic1 promotes septation and that the *fic1-2A* allele may enhance this function.

### *fic1-2A myo2-E1* cells can complete cytokinesis

We next probed this possibility for Fic1 function that would be analogous to Inn1 in *S. cerevisiae* (Sanchez-Diaz et al., 2008). We performed time-lapse imaging at 36°C with wild-type, *fic1-2A, myo2-E1*, and *fic1-2A myo2-E1* cells expressing the membrane marker LactC2-GFP, to monitor membrane ingression, and the SPB marker Sad1-GFP, to monitor mitotic progression (Curto et al., 2014; Hagan and Yanagida, 1995). The kinetics of septation were measured by timing membrane ingression and daughter cell abscission beginning from SPB separation at the onset of mitosis. The timing of anaphase B onset was similar between all genotypes (Fig. 2A,B). The initiation of membrane ingression was similar between wild-type and *fic1-2A* cells, averaging 14.9±0.4 and 16.2±0.3 minutes, respectively (Fig. 2A,C). However, both *myo2-E1* and *fic1-2A myo2-E1* cells exhibited delays in the initiation of membrane ingression compared to wild-type, averaging 28.4±1.0 and 28.8±1.9 minutes, respectively (Fig. 2A,C). Daughter cell separation was completed at similar times in the wild-type and *fic1-2A* cells, averaging 40.6±0.6 and 43.1±0.6 minutes, respectively (Fig. 2A,D). None of the *myo2-E1* daughter cells separated but 7 out of the 22 imaged *fic1-2A myo2-E1* daughter cells took an average time of 117.9±20.2 minutes to separate (Fig. 2A,D). The ability of some *fic1-2A myo2-E1* cells to complete membrane ingression and abscission is consistent with idea that Fic1-2A enhances septum formation.

**Figure 2.**
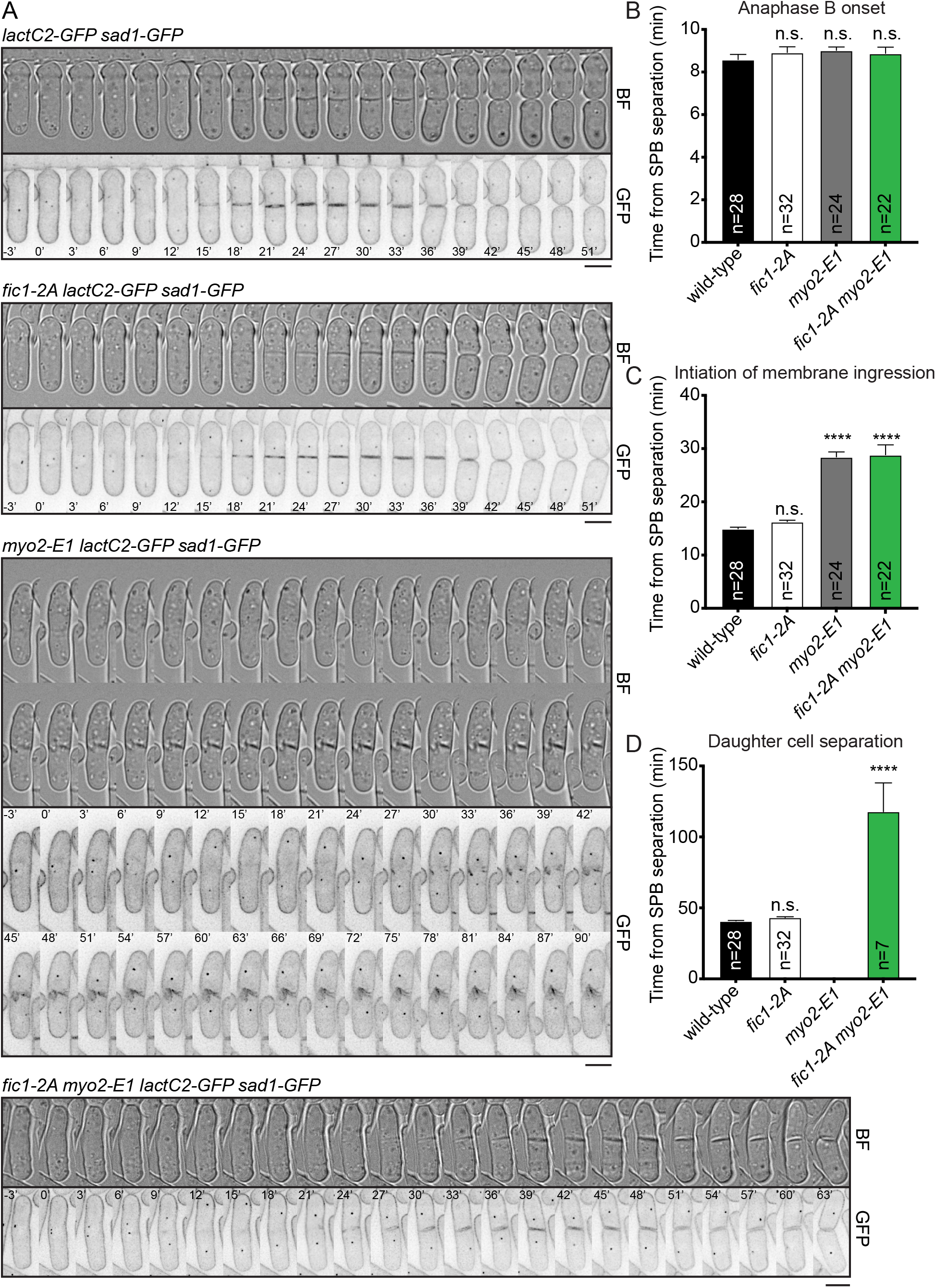
*fic1-2A myo2-E1* cells can achieve membrane ingression and cell separation at *myo2-E1’s* restrictive temperature. A) Representative images of live-cell time-lapse movies from the indicated strains at 36°C. Images were acquired every 3 minutes. Scale bar = 5 μm. B-D) Quantification of timing of anaphase B (B), initiation of membrane ingression (C), and completion of daughter cell separation (D) for each strain. Anaphase B onset was defined as the period from the separation of the SPBs to the initiation of SPB segregation towards opposite cell poles. CR assembly was defined as the period from the separation of the SPBs to the coalescence of cytokinetic nodes into a ring. CR maturation was defined as the period from the completion CR assembly to the initiation of CR contraction. CR constriction was defined as the period from CR contraction to the disappearance of the *rlc1-mNG* from the site of division. n, number of cells analyzed. Data presented as mean ± S.E.M. ****p≤0.0001, n.s., not significant, one-way ANOVA.

### Fic1 directly interacts with Cyk3’s SH3 domain

Because the involvement of Fic1 in promoting septation was reminiscent of the role of *S. cerevisiae*’s IPC (Devrekanli et al., 2012; Nishihama et al., 2009; Sanchez-Diaz et al., 2008), we asked whether Fic1’s interactions with Cdc15 and Imp2 were required for *myo2-E1* suppression. Fic1 binds the SH3 domains of the F-BAR proteins Cdc15 and Imp2 (Roberts-Galbraith et al., 2009) through the P254,257 PxxP motif and the *fic1-P257A* mutation, which disrupts Fic1’s interactions with Cdc15 and Imp2 (Bohnert and Gould, 2012), prevented *fic1-2A*’s suppression of *myo2-E1* (Fig. 3A,B). These data suggest that Fic1’s interaction with Cdc15 and Imp2 are required for *fic1-2A*’s suppression of *myo2-E1* and thus, Fic1’s role in promoting septum formation.

**Figure 3.**
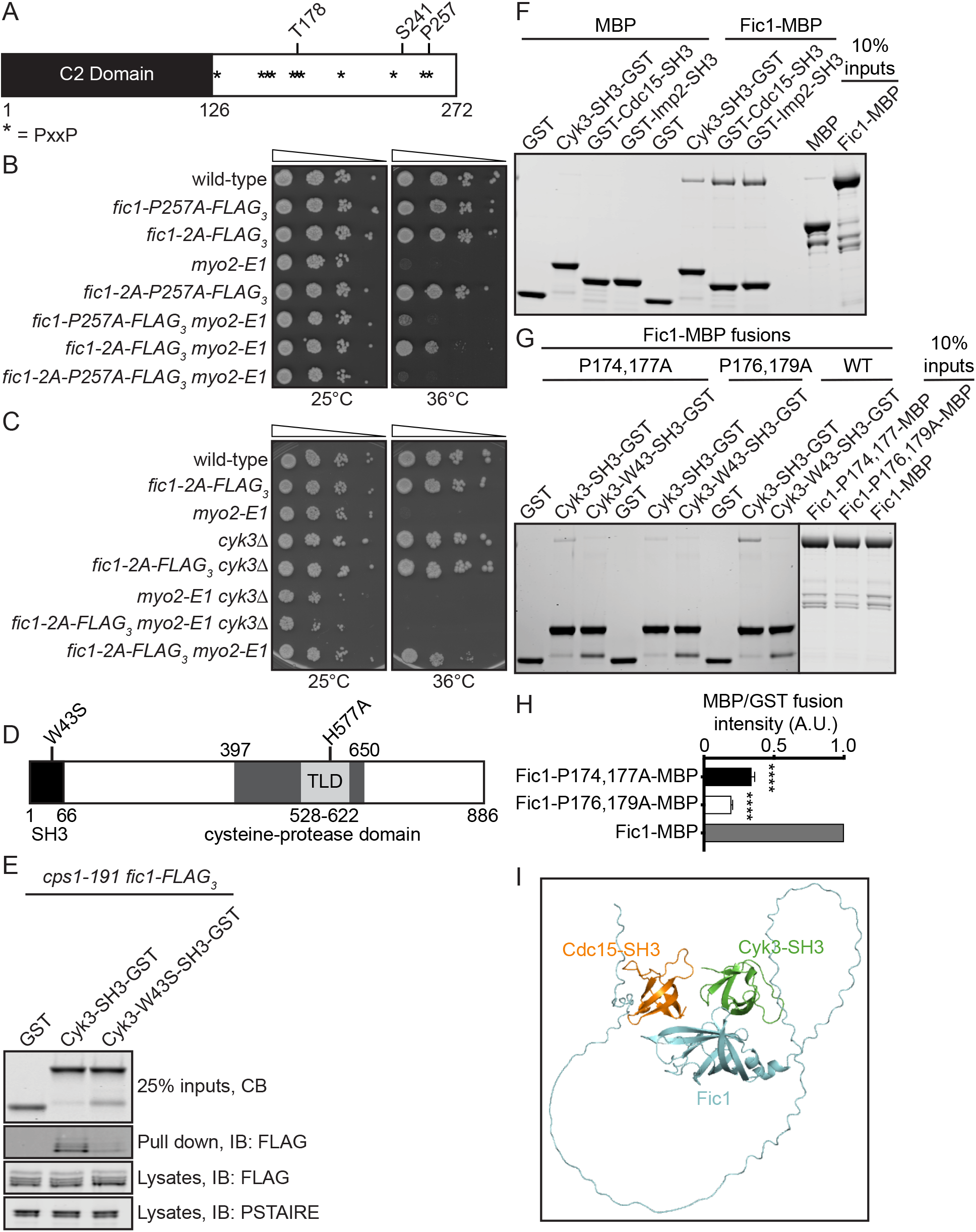
Cyk3-SH3 binds Fic1. A) Schematic of Fic1 with domain boundaries and phosphorylation sites indicated with amino acids numbers and PxxP motifs by asterisks. Drawn to scale. B and C) Ten-fold serial dilutions of the indicated strains were spotted on YE agar media and incubated at the indicated temperatures for 3-5 days. D) Schematic of Cyk3 drawn to scale. Domains and their boundaries and mutations within the domains indicated. E) A portion of protein lysates from *cps1-191 fic1-FLAG*_*3*_ cells was subjected to immunoblotting with FLAG anti-CDK (PSTAIRE) antibodies. The remainder of the lysates was incubated with the indicated bead-bound GST recombinant proteins, of which a portion was detected by Coomassie blue (CB) staining. Fic1 bound to the beads after washing was detected with anti-FLAG immunoblotting. Cells were shifted to 36°C for 3 hours prior to lysis. F and G) Coomassie blue stained SDS–PAGE gel of *in vitro* binding assays using the indicated recombinant proteins. H) Quantification of the amount of soluble protein captured by bead-bound proteins, normalized to the amount of bead-bound protein. Data presented as mean ± S.E.M. ****p≤0.0001, one-way ANOVA. I) Molecular modeling predictions of interactions between Fic1 in cyan, Cyk3-SH3(aa1-66) in green, and Cdc15-SH3(aa867-927) in orange.

We next asked if *S. pombe* Cyk3 was required for *fic1-2A*’s suppression of *myo2-E1*. Indeed, *cyk3Δ* prevented *fic1-*2A from suppressing *myo2-E1* (Fig. 3C). We then aimed to determine if Cyk3 bound Fic1 through an SH3-PxxP interface, similarly to Cyk3 and Inn1 in *S. cerevisiae* (Nishihama et al., 2009) (Fig. 3D). To test this, we generated recombinant Cyk3-SH3-GST and as a negative control, Cyk3-SH3-W43S-GST. Based on SH3 domain homology, the W43S substitution is predicted to disrupt Cyk3-SH3’s ability to bind PxxP motifs (Saksela and Permi, 2012). Immobilized Cyk3-SH3-GST purified Fic1-FLAG_3_ from lysates of *S. pombe* arrested by *cps1-191*, a temperature-sensitive allele of *bgs1* which allows CRs to form but prevents primary septum deposition (Liu et al., 1999), but Cyk3-W43S-SH3-GST did not (Fig. 3E). Finally, we found that Fic1-MBP directly bound Cyk3-SH3-GST indicating that Cyk3-SH3 directly binds Fic1 (Fig. 3F).

To verify that at least one of Fic1’s 11 PxxP motifs was necessary for interaction with Cyk3’s SH3 domain, we generated recombinant Fic1-MBP with every PxxP motif mutated to AxxA, referred to as Fic1-11AxxA-MBP. As predicted, Fic1-11AxxA-MBP did not bind Cyk3-SH3-GST or Cyk3-W43S-SH3-GST (Fig. S2A). To identify which PxxP motif was required for the interaction, we generated Fic1-MBP fusion proteins with each individual AxxA mutation. Fic1-P174,177A-MBP and Fic1-P176,179A-MBP exhibited reduced binding to Cyk3-SH3-GST compared to Fic1-MBP (Fig. 3G,H). Because these are distinct from the PxxP motif involved in binding Cdc15 and Imp2, Fic1 might be able to bind Cyk3 and Cdc15 or Imp2 simultaneously (Bohnert and Gould, 2012) to form an analog of the IPC. Indeed, molecular modeling using ColabFold predicted that Fic1 could simultaneously bind the SH3 domains of Cdc15 and Cyk3 (Fig. 3I and S2B) (Jumper et al., 2021; Mirdita et al., 2022).

Because the Fic1-2A mutant eliminates phosphorylation on T178, we wondered whether disrupting the prolines required for Cyk3 binding around T178 might alter Fic1’s phosphorylation status. We were especially cognizant of this possibility because T178 can be phosphorylated *in vitro* by CDK, a proline-directed kinase (Bohnert et al., 2020). To examine whether these proline mutations affected Fic1 phosphorylation *in vivo*, we analyzed the gel mobilities of Fic1-P174,177A and Fic1-P176,179A. Fic1-FLAG_3_ migrates as four bands. The top band represents dual phosphorylation at T178 and S241, the two intermediate bands are singly phosphorylated at T178 or S241, and the fastest migrating form is not phosphorylated (Bohnert et al., 2020). As predicted, Fic1-P176,179A formed only two bands, consistent with a loss of T178 phosphorylation (Fig. S2C) (Bohnert et al., 2020). Interestingly, Fic1-P174,177A displayed the wild-type pattern of phosphorylation suggesting that it could be used to selectively test the role of Cyk3 binding to Fic1 in the suppression of *myo2-E1* (Fig. S2C). We found that *fic1-2A-P174,177A* did not suppress *myo2-E1* (Fig. S2D) and similarly, inactivation of the Cyk3-SH3 domain by the *cyk3-W43S* allele disrupted *fic1-2A*’s suppression of *myo2-E1* (Fig. S3A). Taken together, these results suggest Cyk3 is required for Fic1’s roles in septum formation.

### Cyk3’s SH3 domain and TLD are required for Fic1’s roles in septum formation

In addition to its SH3 domain, Cyk3 has a central transglutaminase-like domain (TLD) within a larger cysteine protease-like domain (CPD), which has been implicated in Cyk3 function but not thought to have enzymatic activity (Fig. 3D) (Pollard et al., 2012). To determine if Cyk3’s TLD is required for *fic1-2A*’s suppression of *myo2-E1*, we inactivated the TLD through the previously established H577A mutation (Pollard et al., 2012) and found that this mutation also disrupted *fic1-2A*’s suppression of *myo2-E1* (Fig. S3B). To ensure that *cyk3-W43S* and *cyk3-H577A* were not disrupting *fic1-2A*’s suppression by destabilizing Cyk3 or by preventing Cyk3’s localization to the CR, we measured the fluorescence intensity of GFP fusion proteins Cyk3-W43S and Cyk3-H577A. We found that both alleles had similar CR and whole cell fluorescence as Cyk3-GFP (Fig. S3C-E), demonstrating these alleles were stably expressed and localized normally. Additionally, we found that both Cyk3-W43S and Cyk3-H577A co-immunoprecipitated with Cdc15 from prometaphase arrested cells as Cyk3 does (Fig. S3F) (Bohnert and Gould, 2012; Roberts-Galbraith et al., 2010).

### Fic1 and Cyk3 function independently of Chs2

*S. cerevisiae* Cyk3’s TLD stimulates Chs2 (Foltman et al., 2016; Nishihama et al., 2009). While *S. pombe*’s septum lacks chitin, *S. pombe* does have an orthologous protein to *S. cerevisiae*’s Chs2 with the same name but lacking catalytic activity (Horisberger et al., 1978; Martin-Garcia et al., 2003; Matsuo et al., 2004; Sietsma and Wessels, 1990). *S. pombe* Chs2 possibly influences septum formation indirectly because, like *fic1Δ* and *cyk3Δ* cells, *chs2Δ* cells display delays in CR constriction (Martin-Garcia and Valdivieso, 2006). In the cases of *fic1Δ* and *cyk3Δ*, CR constriction delays correlate with a postponement in the onset of bipolar growth, also known as new end take off (NETO), and a transition to invasive pseudophyphal growth (Bohnert and Gould, 2012). If Chs2 acts downstream of Fic1 and Cyk3 we would expect that *chs2Δ* cells to also exhibit NETO defects and invasive pseudophyphal growth. We analyzed bipolar growth establishment in *chs2Δ, fic1Δ, cyk3Δ*, and combination *fic1Δ chs2Δ, cyk3Δ chs2Δ*, and *fic1Δ cyk3Δ chs2Δ* mutants and found that interphase cells of each indicated genotype had an increase in cells growing from only one end (monopolar) compared to wild-type (Fig. 4A). However, *chs2Δ* cells did not exhibit polarity defects at the time of septation or invasive pseudohyphal growth (Fig. 4B,C). Further, deletion of *chs2* did not disrupt *fic1-2A*’s suppression of *myo2-E1* and surprisingly, *chs2Δ* independently suppressed *myo2-E1* (Fig. 4D). These data together indicate that Chs2 does not act downstream of Fic1 and Cyk3 in septation. In accord, ColabFold did not predict an interaction between Cyk3’s CPD and Chs2 in *S. pombe* and even the predicted interaction between Cyk3’s CPD and Chs2 in *S. cerevisiae* was weak (Fig. 4E-H) (Jumper et al., 2021; Mirdita et al., 2022).

**Figure 4.**
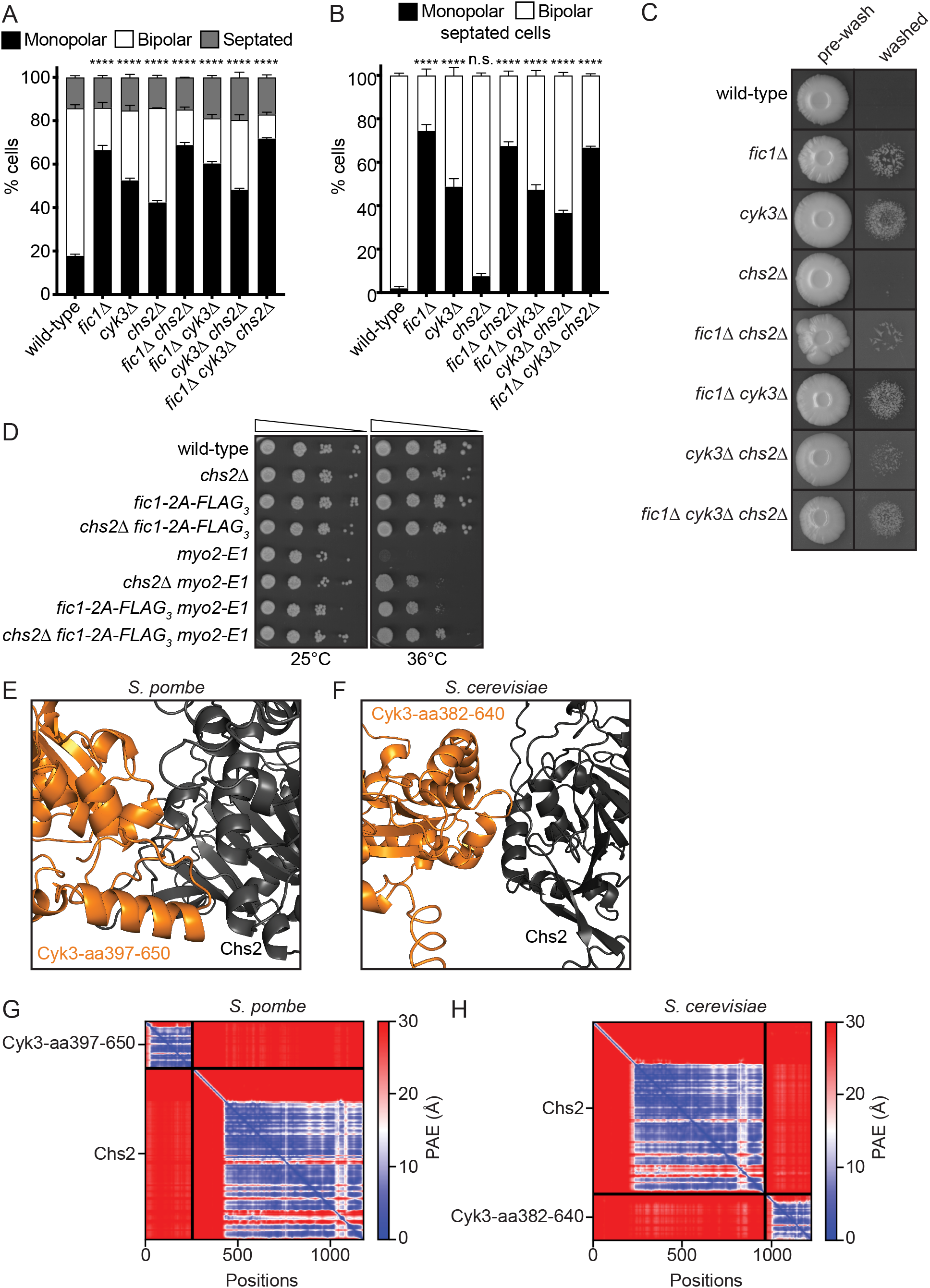
Fic1 functions independently of Chs2. A and B) Quantification of growth polarity phenotypes for interphase (A) and septated cells (B) of the indicated genotypes. Data from three trials per genotype with n > 300 cells for each trial are presented as mean ± S.E.M. The percentage of monopolar cells between wild-type and other genotypes was compared. ****P<0.0001, n.s., not significant, two-way ANOVA with Dunnett’s multiple-comparisons test. C) Invasive growth assays of the indicated genotypes on 2% agar. Cells were spotted on YE agar media and incubated for 20 days at 29°C (top row, “pre-wash”). Colonies were then rinsed under a stream of water and rubbed off (bottom row, “washed”). D) Ten-fold serial dilutions of the indicated strains were spotted on YE agar media and incubated at the indicated temperatures for 3-5 days. E and F) Molecular modeling predictions of the interaction between Cyk3-aa397-650 in orange and Chs2 in grey in *S. pombe* (E) and Cyk3-aa382-640 in orange and Chs2 in grey in *S. cerevisiae* (F). G and H) The predicted aligned error (PAE) map from the molecular modeling of the indicated proteins in *S. pombe* (G) and *S. cerevisiae* (H).

In conclusion, our results suggest that mutating the two Fic1 phosphorylation sites to alanine enhances Fic1’s normal role in septation to allow *myo2-E1* suppression. Whether this is due to preventing Fic1 phosphorylation or changing Fic1 structure remains to be determined. However, Fic1’s interactions with Cyk3, Cdc15 and/or Imp2 are required for this function and molecular modeling suggests Fic1 could simultaneously bind the SH3 domains of Cyk3 and Cdc15/Imp2 to form a complex similar to the *S. cerevisiae* IPC. Despite similar interactions to *S. cerevisiae*’s IPC, these proteins promote septum formation independent of Chs2 and it will be interesting to determine how they influence this critical aspect of fission yeast cell division.

## Abbreviations

(CR): cytokinetic ring
(NETO): new end take off
(SPB): spindle pole body
(TLD): transglutaminase-like domain
(IPC): ingression progression complex
(CPD): cysteine-protease domain
(PAE): predicted aligned error

## Acknowledgements

We are grateful to Yolanda Sánchez for providing strains. We thank Alaina Willet, Sierra Cullati, Kazutoshi Akizuki, and Jun-Song Chen for critical reading of the manuscript.

## Materials and Methods

### Yeast methods

*Schizosaccharomyces pombe* strains utilized in this study (Supplemental Table S1) were cultured in yeast extract (YE) media (Moreno et al., 1991). Glutamate media was used for crosses (Moreno et al., 1991). *fic1, cyk3, sid4, rlc1*, and *sad1* were tagged endogenously at the 3′ end of their open reading frames (ORFs) with *FLAG*_*3*_*:kan*^*R*^, *GFP:kan*^*R*^, *mNG:hyg*^*R*^, *V5*_*3*_*:hyg*^*R*^, and/or *mCherry:nat*^*R*^ using pFA6 cassettes as previously described (Bahler et al., 1998). G418 (100 μg/ml; Sigma-Aldrich, St. Louis, MO), Hygromycin B (125 μg/mL; Invitrogen, Waltham, MA), and Nourseothricin (125 μg/mL; Gold Biotechnology St. Louis, MO) in YE media was used for selecting *kan*^*R*^, *hyg*^*R*^, and *nat*^*R*^ cells respectively. *mNG*, a YFP derivative from the lancelet *Branchiostoma lanceolatum*, was selected for imaging experiments because of its superior brightness (Shaner et al., 2013; Willet et al., 2015). A lithium acetate transformation method (Keeney and Boeke, 1994) was used for introducing tagging sequences, and endogenous integration of tags were verified by whole-cell PCR and/or microscopy. Introduction of tagged loci into other genetic backgrounds was accomplished using standard *S. pombe* mating, sporulation, and tetrad-dissection techniques. Fusion proteins were expressed from their endogenous locus under control of their native promoter unless otherwise indicated. For serial-dilution growth assays, cells were cultured in liquid YE at 25°C in a shaking incubator, four 1:10 serial dilutions starting at 1.5 × 10^6^ cells/mL were made, 2 μL of each dilution was spotted on YE agar, and cells were grown at the specified temperatures for 3-5 days. All spot assays were performed in triplicate and representative images are shown.

Mutants of *fic1* were expressed from the endogenous loci. To make *fic1* mutations, *fic1*^*+*^ gDNA with 500 bp 5’ and 3’ flanks was inserted between BamHI and PstI sites of pIRT2 (Bohnert and Gould, 2012) and site-directed mutagenesis was used to introduced the desired mutations, which were confirmed by DNA sequencing. *fic1::ura*^*+*^ was transformed with these pIRT2-*fic1* constructs, and stable integrants resistant to 1.5 g/L 5-fluoroorotic acid (5-FOA) (Fisher Scientific, Hampton, NH) were isolated. The correct insertion site and mutations were confirmed by whole-cell PCR and DNA sequencing.

*cyk3* strains were made in a similar manner but pIRT2-*cyk3* (W43S, H577A, W43S,H577A) mutant plasmids were constructed from *cyk3* cDNA with 500 bp 5’ and 3’ flanks to allow these *cyk3* alleles to be verified by whole-cell PCR once integrated.

### Invasive growth assays

To assay pseudohyphal invasive growth, 5 μL containing a total of 10^5^ cells were spotted on 2% YE agar and incubated at 29°C for 20 days. Colonies were subsequently placed under a steady stream of water and surface growth was wiped off using a paper towel, as described previously (Pohlmann and Fleig, 2010; Prevorovsky et al., 2009).

### Microscopy

Live-cell imaging of *S. pombe* cells were acquired using one of the following: (1) a personal DeltaVision microscope system (Leica Microsystems, Wetzlar, Germany) that includes an Olympus IX71 microscope, 60× NA 1.42 PlanApo and 100× NA 1.40 UPlanSApo objectives, a pco.edge 4.2 sCMOS camera, and softWoRx imaging software or (2) a Zeiss Axio Observer inverted epifluorescence microscope with Zeiss 63X Oil (1.46 NA), a Axiocam 503 monochrome camera (Zeiss), and captured using Zeiss ZEN 3.0 (Blue edition) software. All cells were in log phase growth before temperature-sensitive shifts and/or live imaging. Time-lapse imaging was performed using an ONIX microfluidics perfusion system (CellASIC ONIX; EMD Millipore, Burlington, MA). A suspension of 50 μL of 40 × 10^6^ cells/ml YE was loaded into Y04C plates for 5 s at 8 psi. YE media was flowed through the chamber at 5 psi throughout imaging.

Anaphase B onset was defined as the period from the separation of the SPBs to the initiation of SPB segregation towards opposite cell poles. CR assembly was defined as the period from the separation of the SPBs to the coalescence of the cytokinetic nodes into a clearly defined ring. CR maturation was defined as the period from the completion CR assembly to the initiation of CR contraction. CR constriction was defined as the period from CR contraction to the disappearance of the *rlc1-mNG* from the site of division.

Intensity measurements were made from non-deconvolved summed Z-projections of the images processed through ImageJ software (Schneider et al., 2012). For all intensity measurements, the background was subtracted by selecting a region of interest (ROI) in the same image in an area free of cells. The background raw intensity was divided by the area of the background, which was multiplied by the area of the measured object. This number was then subtracted from the intensity measurement of that object. Max intensity Z projections are shown in representative images.

To visualize birth scars by Calcofluor staining, cells were washed in PBS and then resuspended in PBS containing 5 μg/mL Calcofluor and allowed to incubate on ice for 30 minutes. Cells were then washed three times in PBS and images were acquired. Using the proximity of birth scars to cell ends, growth/morphology was categorized as one of the following: monopolar (i.e. growth on one end), bipolar (i.e. growth on both ends), monopolar and septated, or bipolar and septated. All cells stained with Calcofluor were grown to log phase at 25°C.

### Protein Methods

Cells were lysed by bead disruption in NP-40 buffer in denaturing conditions as previously described (Gould et al., 1991). Immunoblot analysis of cell lysates and immunoprecipitates was performed using anti-FLAG (M2; Sigma-Aldrich, St. Louis, MO), anti-PSTAIR Cdc2 (Sigma-Aldrich, St, Louis, MO), anti-GFP (Roche, Indianapolis, IN), or anti-GFP (VUIC9H4) antibodies or serums raised against GST-Cdc15 (amino acids 1–405; VU326; Cocalico Biologicals, Stevens, PA) as previously described (Bohnert et al., 2009). Cyk3-SH3-GST (aa1-66), Fic1-MBP, GST-Cdc15-SH3 (aa867-927), and GST-Imp2-SH3 (aa608-670) recombinant proteins and their variants were purified from *E. coli* using standard biochemistry techniques. *in vitro* binding assays were performed in 20 mM Tris (pH 7.4) and 150 mM NaCl and allowed to incubated at 4°C for 1 hour while nutating. The beads were washed 3 times with reaction buffer before peforming SDS-PAGE and Coomassie staining.

Affinity purifications using bead bound Cyk3-SH3-GST recombinant proteins and *S. pombe* lysate were performed by lysing cells in Cyk3 lysis buffer (50 mM Tris (pH 7.5), 150 mM NaCl, 1 mM EDTA, 50 mM NaF, 0.5% NP-40, 0.1% SDS, 1 mM DTT, 1 mM PMSF, 1.3 mM Benzamidine, and cOmplete protease inhibitors (Roche, Indianapolis, IN). *S. pombe* lysates and Cyk3-SH3-GST recombinant proteins were incubated at 4°C for 1 hour while nutating. The bead bound recombinant proteins were washed 3 times in the lysate buffer before SDS-PAGE and immunoblotting or Coomassie staining.

## Supplemental Figure Legends

**Figure S1.**
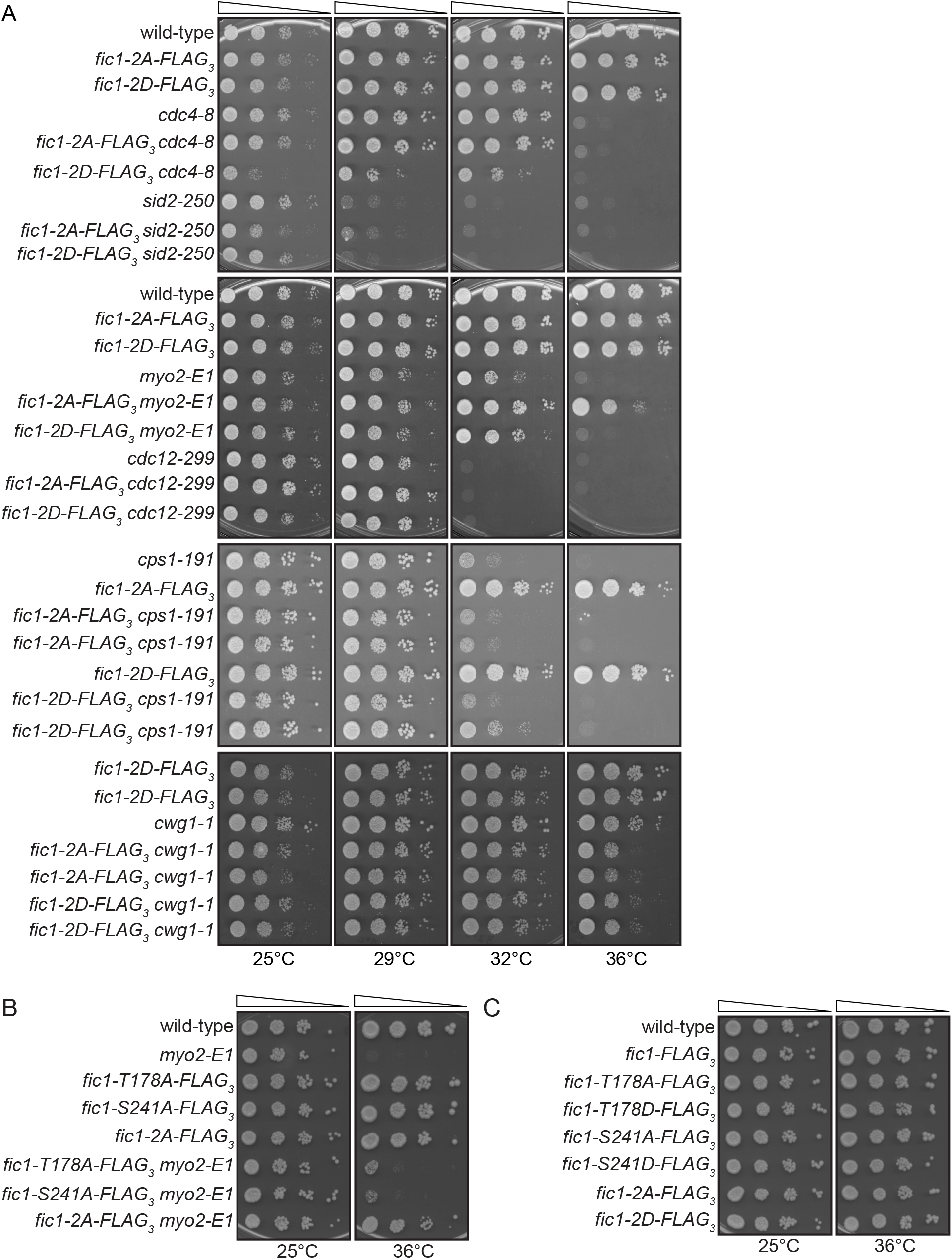
Individual or single *fic1* phosphomutants are not temperature-sensitive. A-C) Ten-fold serial dilutions of the indicated strains were spotted on YE agar media and incubated at the indicated temperatures for 3-5 days.

**Figure S2.**
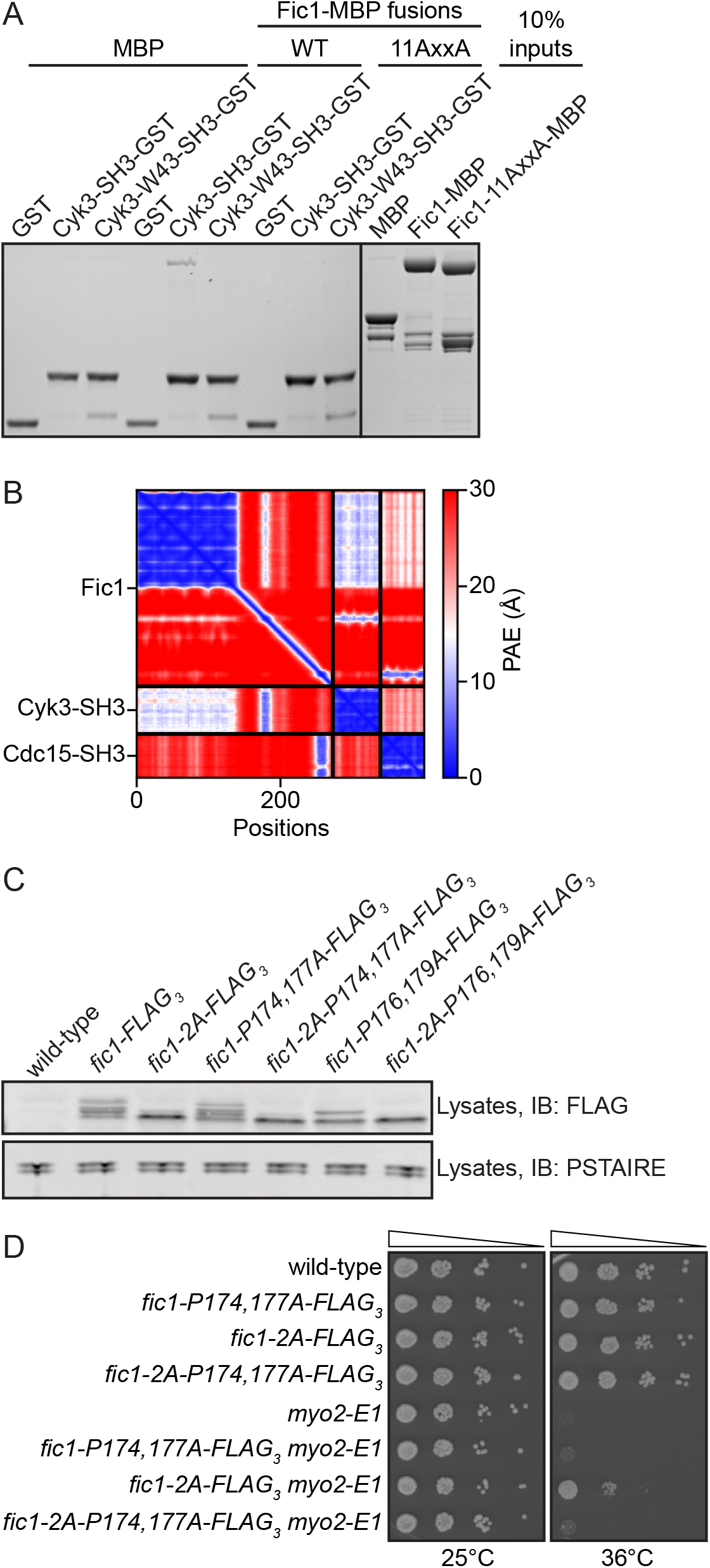
P174,177 is required for *fic1-2A*’s suppression of *myo2-E1*. A) Coomassie-stained SDS–PAGE of *in vitro* binding assays using the indicated recombinant proteins. B) The predicted aligned error (PAE) map from the molecular modeling between Fic1, Cyk3-SH3, and Cdc15-SH3. C) Lysates from cells of the indicated genotypes were immunoblotted with anti-FLAG antibody to assess Fic1-FLAG_3_ gel mobilities and anti-CDK (PSTAIRE) antibody as a loading control. D) Ten-fold serial dilutions of the indicated strains were spotted on YE agar media and incubated at the indicated temperatures for 3-5 days.

**Figure S3.**
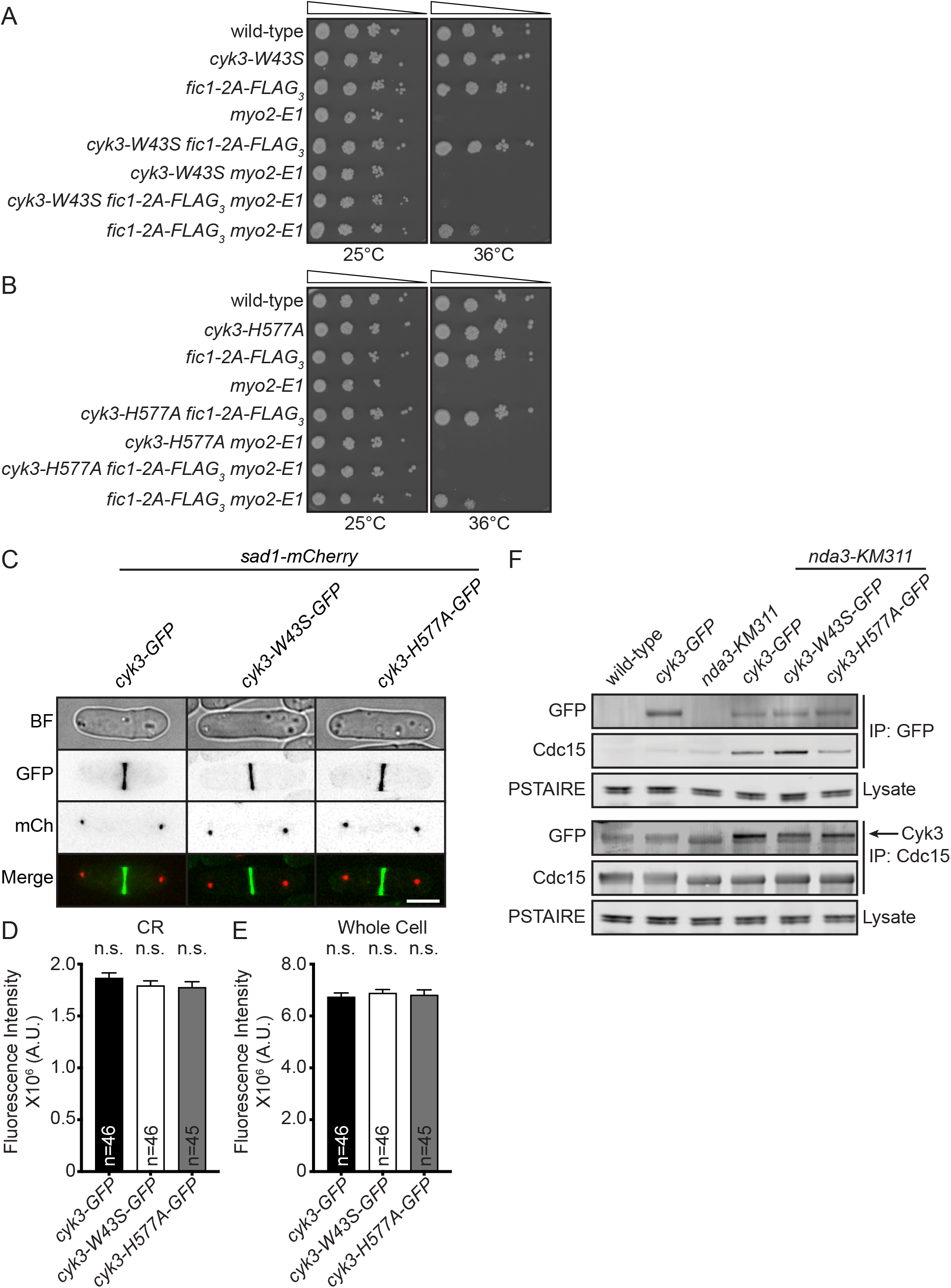
Cyk3’s SH3 and transglutaminase-like domain are required for *fic1-2A’s* suppression of *myo2-E1*. A and B) Ten-fold serial dilutions of the indicated strains were spotted on YE agar media and incubated at the indicated temperatures for 3-5 days. C) Live-cell bright field (BF), GFP, mCherry (mCh) and merged GFP/mCh images of cells of indicated genotypes during cytokinesis. Scale bar: 5 μm. D and E) Quantification of CR (D) and whole cell (E) fluorescence intensities for cells of indicated genotypes. Data from three trials per genotype presented as mean ± S.E.M. n.s., not significant, one-way ANOVA. F) Anti-GFP or anti-Cdc15 immunoprecipitates from cells of indicated genotypes were blotted with an anti-GFP or anti-Cdc15 antibody. Lysate samples were blotted with anti-CDK (PSTAIRE) as an input control for the immunoprecipitations. Arrow indicates Cyk3-GFP protein band.

**TABLE S1.** *S. pombe* strains used in this study.

|  |  |  |
| --- | --- | --- |
| **Figure 1** |  |  |
| KGY246 | *ade6-M210 ura4-D18 leu1-32 h<sup>-</sup>* | Lab Stock |
| KGY11856 | *fic1-T178A,S241A-FLAG<sub>3</sub>:kan<sup>R</sup> ade6-M210 ura4-D18 leu1-32 h<sup>+</sup>* | Lab Stock |
| KGY11861 | *fic1-T178D,S241D-FLAG<sub>3</sub>:kan<sup>R</sup> ade6-M210 ura4-D18 leu1-32 h<sup>+</sup>* | Lab Stock |
| KGY2971 | *myo2-E1 ade6-M216 leu1-32 ura4-D18 h<sup>+</sup>* | Lab Stock |
| KGY1018-2 | *myo2-E1 fic1-T178A,S241A-FLAG<sub>3</sub>:kan<sup>R</sup> ade6-M210 leu1-32 ura4-D18 h<sup>-</sup>* | This Study |
| KGY15368 | *myo2-E1 fic1-T178D,S241D-FLAG<sub>3</sub>:kan<sup>R</sup> ade6-M210 leu1-32 ura4-D18 h<sup>+</sup>* | This Study |
| KGY6008 | *fic1::ura4<sup>+</sup> leu1-32 ura4-D18 ade6-M21X h<sup>+</sup>* | Lab Stock |
| KGY6665 | *fic1::ura4<sup>+</sup> myo2-E1 leu1-32 ura4-D18 ade6-M21X h<sup>+</sup>* | Lab Stock |
| KGY5305-2 | *sid4-GFP:kan<sup>R</sup> rlc1-mNeonGreen:hyg<sup>R</sup> ade6-M210 ura4-D18 leu1-32 h<sup>+</sup>* | This Study |
| KGY5659-2 | *fic1::ura4<sup>+</sup> sid4-GFP:kan<sup>R</sup> rlc1-mNeonGreen:hyg<sup>R</sup> ade6-M21X ura4-D18 leu1-32 h<sup>-</sup>* | This Study |
| KGY5602-2 | *fic1-T178A,S241A-FLAG<sub>3</sub>:kan<sup>R</sup> sid4-GFP:kan<sup>R</sup> rlc1-mNeonGreen:hyg<sup>R</sup> ade6-M210 ura4-D18 leu1-32 h<sup>+</sup>* | This Study |
| KGY5334-2 | *fic1-T178D,S241D-FLAG<sub>3</sub>:kan<sup>R</sup> sid4-GFP:kan<sup>R</sup> rlc1-mNeonGreen:hyg<sup>R</sup> ade6-M210 ura4-D18 leu1-32 h<sup>+</sup>* | This Study |
| **Figure 2** |  |  |
| KGY2152-2 | *act1:LactC2-GFP:nat<sup>R</sup> sad1-GFP:kan<sup>R</sup> ade6-M210 ura4-D18 leu1-32 h<sup>+</sup>* | (Curto et al., 2014) |
| KGY4102-2 | *fic1-T178A,S241A-FLAG<sub>3</sub>:kan<sup>R</sup> act1:LactC2-GFP:nat<sup>R</sup> sad1-GFP:kan<sup>R</sup> ade6-M210 ura4-D18 leu1-32 h<sup>+</sup>* | This Study |
| KGY3655-2 | *myo2-E1 act1:LactC2-GFP:nat<sup>R</sup> sad1-GFP:kan<sup>R</sup> ade6-M210 ura4-D18 leu1-32 h<sup>+</sup>* | This Study |
| KGY4138-2 | *fic1-T178A,S241A-FLAG<sub>3</sub>:kan<sup>R</sup> myo2-E1 act1:LactC2-GFP:nat<sup>R</sup> sad1-GFP:kan<sup>R</sup> ade6-M210 ura4-D18 leu1-32 h<sup>+</sup>* | This Study |
| **Figure 3** |  |  |
| KGY11876 | *fic1-P257A-FLAG<sub>3</sub>:kan<sup>R</sup> leu1-32 ura4-D18 ade6-M21X h<sup>+</sup>* | Lab stock |
| KGY11934 | *fic1-T178A,S241A,P257A-FLAG<sub>3</sub>:kan<sup>R</sup> leu1-32 ura4-D18 ade6-M21X h<sup>+</sup>* | This Study |
| KGY3373-2 | *fic1-P257A-FLAG<sub>3</sub>:kan<sup>R</sup> myo2-E1 leu1-32 ura4-D18 ade6-M21X h<sup>-</sup>* | This Study |
| KGY19546 | *fic1-T178A,S241A,P257A-FLAG<sub>3</sub>:kan<sup>R</sup> myo2-E1 leu1-32 ura4-D18 ade6-M21X h<sup>-</sup>* | This Study |
| KGY7490 | *cyk3::ura<sup>+</sup> ade6-M21X leu1-32 ura4-D18 h<sup>-</sup>* | Lab stock |
| KGY15446 | *cyk3::ura4<sup>+</sup> fic1-T178,S241A-FLAG<sub>3</sub>:kan<sup>R</sup> ade6-M21X leu1-32 ura4-D18 h<sup>+</sup>* | This Study |
| KGY7635 | *cyk3::ura4<sup>+</sup> myo2-E1 ade6-M21X leu1-32 ura4-D18 h<sup>-</sup>* | This Study |
| KGY15443 | *cyk3::ura4<sup>+</sup> fic1-T178,S241A-FLAG<sub>3</sub>:kan<sup>R</sup> myo2-E1 ade6-M21X leu1-32 ura4-D18 h<sup>+</sup>* | This Study |
| KGY3873-2 | *cps1-191 fic1-FLAG<sub>3</sub>:kan<sup>R</sup> chs2-V5<sub>3</sub>:hyg<sup>R</sup> lys1-131 ade6-M21X ura4-D18 leu1-32 h<sup>-</sup>* | This Study |
| **Figure 4** |  |  |
| KGY6365-2 | *chs2::kan<sup>R</sup> ade6-M210 ura4-D18 leu1-32 h<sup>-</sup>* | This Study |
| KGY6580-2 | *chs2::kan<sup>R</sup> fic1-T178A,S241A-FLAG<sub>3</sub>:kan<sup>R</sup> ade6-M210 ura4-D18 leu1-32 h<sup>-</sup>* | This Study |
| KGY6570-2 | *chs2::kan<sup>R</sup> myo2-E1 ade6-M210 ura4-D18 leu1-32 h<sup>+</sup>* | This Study |
| KGY6571-2 | *chs2::kan<sup>R</sup> fic1-T178A,S241A-FLAG<sub>3</sub>:kan<sup>R</sup> myo2-E1 ade6-M210 ura4-D18 leu1-32 h<sup>+</sup>* | This Study |
| KGY6434-2 | *fic1::ura4<sup>+</sup> chs2::kan<sup>R</sup> leu1-32 ura4-D18 ade6-M21X h<sup>+</sup>* | This Study |
| KGY15954 | *fic1::ura4<sup>+</sup> cyk3::ura4<sup>+</sup> leu1-32 ura4-D18 ade6-M21X h<sup>-</sup>* | Lab stock |
| KGY6930-2 | *chs2::kan<sup>R</sup> cyk3::ura4<sup>+</sup> ade6-M21X leu1-32 ura4-D18 h<sup>-</sup>* | This Study |
| KGY7101-2 | *chs2::kan<sup>R</sup> cyk3::ura4<sup>+</sup> fic1::ura4<sup>+</sup> ade6-M21X leu1-32 ura4-D18 h<sup>-</sup>* | This Study |
| **Supplemental Figures** |  |  |
| **Figure S1** |  |  |
| KGY441 | *cdc4-8 ade6-M210 ura4-D18 h<sup>+</sup>* | Lab Stock |
| KGY15363 | *fic1-T178A,S241A-FLAG<sub>3</sub>:kan<sup>R</sup> cdc4-8 ade6-M21X ura4-D18 h<sup>-</sup>* | This Stock |
| KGY15364 | *fic1-T178A,S241D-FLAG<sub>3</sub>:kan<sup>R</sup> cdc4-8 ade6-M21X ura4-D18 h<sup>+</sup>* | This Stock |
| KGY1105 | *sid2-250 ade6-M21X ura4-D18 leu1-32 h<sup>-</sup>* | Lab Stock |
| KGY15374 | *fic1-T178,S241A-FLAG<sub>3</sub>:kan<sup>R</sup> sid2-250 ade6-M210 ura4-D18 leu1-32 h<sup>-</sup>* | This Study |
| KGY15375 | *fic1-T178,S241D-FLAG<sub>3</sub>:kan<sup>R</sup> sid2-250 ade6-M210 ura4-D18 leu1-32 h<sup>-</sup>* | This Study |
| KGY748 | *cdc12-299 h<sup>-</sup>* | Lab Stock |
| KGY15365 | *fic1-T178,S241A-FLAG<sub>3</sub>:kan<sup>R</sup> cdc12-299 leu1-32 ura4-D18 h<sup>+</sup>* | This Study |
| KGY15366 | *fic1-T178,S241D-FLAG<sub>3</sub>:kan<sup>R</sup> cdc12-299 299 leu1-32 ura4-D18 h<sup>+</sup>* | This Study |
| KGY17207 | *cps1-191 ura4-D18 leu1-32 ade6-M210 lys1-131 h<sup>+</sup>* | Lab Stock |
| KGY17771 | *cps1-191 fic1-T178A,S241A-FLAG<sub>3</sub>:kan<sup>R</sup> ura4-D18 ade6-M21X h<sup>-</sup>* | This Study |
| KGY17773 | *cps1-191 fic1-T178D,S241D-FLAG<sub>3</sub>:kan<sup>R</sup> ura4-D18 ade6-M21X h<sup>-</sup>* | This Study |
| KGY11106 | *cwg1-1 leu1-32 h<sup>-</sup>* | Lab Stock |
| KGY17681 | *cwg1-1 fic1-T178A,S241A-FLAG<sub>3</sub>:kan<sup>R</sup> ade6-M21X ura4-D18 leu1-32 h<sup>-</sup>* | This Study |
| KGY17683 | *cwg1-1 (bgs4 ts) fic1-T178D,S241D-FLAG<sub>3</sub>:kan<sup>R</sup> ade6-M21X ura4-D18 leu1-32 h<sup>-</sup>* | This Study |
| KGY19522 | *fic1-T178A-FLAG<sub>3</sub>:kan<sup>R</sup> ade6-M210 ura4-D18 leu1-32 h<sup>+</sup>* | Lab Stock |
| KGY19523 | *fic1-S241A-FLAG<sub>3</sub>:kan<sup>R</sup> ade6-M210 ura4-D18 leu1-32 h<sup>+</sup>* | Lab Stock |
| KGY7615-2 | *fic1-T178A-FLAG<sub>3</sub>:kan<sup>R</sup> myo2-E1 ade6-M210 ura4-D18 leu1-32 h<sup>-</sup>* | This Study |
| KGY7628-2 | *fic1-S241A-FLAG<sub>3</sub>:kan<sup>R</sup> myo2-E1 ade6-M210 ura4-D18 leu1-32 h<sup>-</sup>* | This Study |
| KGY6288 | *fic1-FLAG<sub>3</sub>:kan<sup>R</sup> ade6-M216 leu1-32 ura4-D18 h<sup>-</sup>* | Lab stock |
| KGY11908 | *fic1-T178D-FLAG<sub>3</sub>:kan<sup>R</sup> ade6-M210 ura4-D18 leu1-32 h<sup>+</sup>* | Lab stock |
| KGY11854 | *fic1-S241D-FLAG<sub>3</sub>:kan<sup>R</sup> leu1-32 ura4-D18 ade6-M21X h<sup>+</sup>* | Lab stock |
| **Figure S2** |  |  |
| KGY4705-2 | *fic1-P174,177A-FLAG<sub>3</sub>:kan<sup>R</sup> leu1-32 ura4-D18 ade6-M21X h<sup>+</sup>* | This Study |
| KGY4774-2 | *fic1-T178A,S241A-P174,177A-FLAG<sub>3</sub>:kan<sup>R</sup> leu1-32 ura4-D18 ade6-M21X h<sup>+</sup>* | This Study |
| KGY4706-2 | *fic1-P176,179A-FLAG<sub>3</sub>:kan<sup>R</sup> leu1-32 ura4-D18 ade6-M21X h<sup>+</sup>* | This Study |
| KGY4708-2 | *fic1-T178A,S241A-P176,179A-FLAG<sub>3</sub>:kan<sup>R</sup> leu1-32 ura4-D18 ade6-M21X h<sup>+</sup>* | This Study |
| KGY4717-2 | *fic1-P174,177A-FLAG<sub>3</sub>:kan<sup>R</sup> myo2-E1 leu1-32 ura4-D18 ade6-M21X h<sup>+</sup>* | This Study |
| KGY4852-2 | *fic1-T178A,S241A-P174,177A-FLAG<sub>3</sub>:kan<sup>R</sup> myo2-E1 leu1-32 ura4-D18 ade6-M21X h<sup>-</sup>* | This Study |
| **Figure S3** |  |  |
| KGY899-2 | *cyk3-W43S ade6-M21X leu1-32 ura4-D18 h<sup>-</sup>* | This Study |
| KGY2086-2 | *cyk3-W43S fic1-T178A,S241A-FLAG<sub>3</sub>:kan<sup>R</sup> ade6-M21X leu1-32 ura4-D18 h<sup>+</sup>* | This Study |
| KGY2141-2 | *cyk3-W43S myo2-E1 ade6-M21X leu1-32 ura4-D18 h<sup>-</sup>* | This Study |
| KGY2505-3 | *cyk3-W43S fic1-T178A,S241A-FLAG<sub>3</sub>:kan<sup>R</sup> myo2-E1 ade6-M21X leu1-32 ura4-D18 h<sup>-</sup>* | This Study |
| KGY1886-2 | *cyk3-H577A ade6-M21X leu1-32 ura4-D18 h<sup>-</sup>* | This Study |
| KGY2120-2 | *cyk3-H577A fic1-T178A,S241A-FLAG<sub>3</sub>:kan<sup>R</sup> ade6-M21X leu1-32 ura4-D18 h<sup>+</sup>* | This Study |
| KGY2161-2 | *cyk3-H577A myo2-E1 ade6-M21X leu1-32 ura4-D18 h<sup>-</sup>* | This Study |
| KGY2559-2 | *cyk3-H577A fic1-T178A,S241A-FLAG<sub>3</sub>:kan<sup>R</sup> myo2-E1 ade6-M21X leu1-32 ura4-D18 h<sup>-</sup>* | This Study |
| KGY6897 | *cyk3-FLAG<sub>3</sub>:kan<sup>R</sup> ade6-M21X leu1-32 ura4-D18 h<sup>-</sup>* | Lab Stock |
| KGY2460-2 | *cyk3-GFP:kan<sup>R</sup> sad1-mCherry:nat<sup>R</sup> ade6-M210 ura4-D18 leu1-32 h<sup>-</sup>* | This Study |
| KGY2362-2 | *cyk3-W43S-GFP:kan<sup>R</sup> sad1-mCherry:nat<sup>R</sup> ade6-M21X leu1-32 ura4-D18 h<sup>-</sup>* | This Study |
| KGY2364-2 | *cyk3-H577A-GFP:kan<sup>R</sup> sad1-mCherry:nat<sup>R</sup> ade6-M21X leu1-32 ura4-D18 h<sup>-</sup>* | This Study |
| KGY2202-3 | *cyk3-GFP:kan<sup>R</sup> ade6-M210 ura4-D18 leu1-32 h<sup>-</sup>* | Lab Stock |
| KGY56 | *nda3-km311 leu1-32 h<sup>+</sup>* | Lab Stock |
| KGY2890-2 | *cyk3-GFP:kan<sup>R</sup> nda3-km311 ade6-M210 ura4-D18 leu1-32 h<sup>-</sup>* | This Study |
| KGY2895-2 | *cyk3-W43S-GFP:kan<sup>R</sup> nda3-km311 ura4-D18 leu1-32 h<sup>-</sup>* | This Study |
| KGY2892-2 | *cyk3-H577A-GFP:kan<sup>R</sup> nda3-km311 ade6-M210 ura4-D18 leu1-32 h<sup>-</sup>* | This Study |

